# Field evaluation of a quantitative, and rapid malaria diagnostic system using a fluorescent Blue-ray optical device

**DOI:** 10.1101/721076

**Authors:** Takeki Yamamoto, Muneaki Hashimoto, Kenji Nagatomi, Takahiro Nogami, Yasuyuki Sofue, Takuya Hayashi, Yusuke Ido, Shouki Yatsushiro, Kaori Abe, Kazuaki Kajimoto, Noriko Tamari, Beatrice Awuor, George Sonye, James Kongere, Stephen Munga, Jun Ohashi, Hiroaki Oka, Noboru Minakawa, Masatoshi Kataoka, Toshihiro Mita

## Abstract

We improved a previously developed quantitative malaria diagnostic system based on fluorescent Blue-ray optical device. Here, we first improved the diagnostic system to enable fully automated operation and the field application was evaluated in Kenya. We detected *Plasmodium falciparum* in blood samples collected from 288 individuals aged 1-16 years using nested polymerase chain reaction (nPCR), rapid diagnostic test (RDT), and automated system. Compared to RDT, the automated system exhibited a higher sensitivity (100%; 95% confidence interval [CI], 93.3–100%) and specificity (92.8%; 95%CI, 88.5–95.8%). The limit of detection was 0.0061%. Linear regression analysis revealed a correlation between the automated system and microscopic examination for detecting parasitemia (adjusted R^2^ value=0.63, P=1.13×10^−12^). The automated system exhibited a stable quantification of parasitemia and a higher diagnostic accuracy for parasitemia than RDT. This indicates the potential of this system as a valid alternative to conventional methods used at local health facilities, which lack basic infrastructure.

## Introduction

Globally, malaria is one of the “big three” infectious diseases with approximately 37 million people at risk for contracting malaria (WHO, 2018). Malaria is a vector-borne disease, which is caused by infection from *Plasmodium* spp. and is transmitted by *Anopheles* mosquitoes. The United Nations Sustainable Development Goals had proposed to end the epidemic of malaria by 2030 (WHO, 2015). Although the annual case fatality rate has not increased recently, 435,000 malaria-related deaths were recorded in over 100 countries in 2017 (WHO, 2018). Since 2014, new malaria cases have increased slightly with 219 million recorded cases in 2017 (Alonso and Noor, 2017; WHO, 2018). Several factors have stagnated the global progress against malaria, such as the emergence of insecticide-resistant vectors, poor access to insecticide-treated nets and artemisinin-based combination therapies, and inefficiency of tools that are currently used in malaria diagnosis, treatment, and control (Alonso and Noor, 2017). Particularly, most tools were developed before the year 2000 and are not efficient to tackle the current malaria infection. Therefore, novel technologies must be scaled up to accelerate the development of new tools for malarial control and eradication (Alonso and Noor, 2017).

Malaria must be accurately diagnosed to ensure effective patient management and to prevent unnecessary treatment. Rapid diagnostic tests (RDTs), which detect malaria-specific antigens, such as histidine-rich protein 2 (HRP2), are easy to use, rapid, and affordable. Hence, RDTs are widely used even in remote areas, which do not have access to microscopic diagnosis. RDTs have wide sensitivity (63–100%) and specificity (53–100%) ranges for detecting *Plasmodium falciparum* (Boyce and O’Meara, 2017). However, RDTs are unable to determine the parasite infection rate in the red blood cells (RBCs) of peripheral blood (parasitemia), which can lead to the misdiagnosis of severe malaria. Additionally, the clearance of HRP2 antigen in the patient’s blood is very slow. Therefore, the presence of residual antigen potentially produces a persistent malaria-positive status even after several weeks post parasite clearance (Aydin-Schmidt et al., 2013). Furthermore, blood samples containing *Plasmodium falciparum* with deleted *HRP2* gene can be diagnosed as false-negative in RDT (Parr et al., 2016). Microscopic examination of Giemsa-stained blood smear is a low-cost method for diagnosing parasitemia and for differentiating *Plasmodium* species. However, microscopic examination involves labor-intensive steps and requires technical expertise for accurate diagnosis. Additionally, numerous studies suggest that submicroscopic malaria infections, which are not detectable by microscopy, can be a source of malaria transmission from humans to mosquitoes (Bousema et al., 2012; Bousema et al., 2014; Goncalves et al., 2017; Lin et al., 2015; Okell et al., 2012; Ouedraogo et al., 2009; Tadesse et al., 2018). The currently available malaria diagnostic methods cannot detect submicroscopic infections.

Recently, we had developed a portable, easy to operate, and battery driven fluorescent Blue-ray optical device for determining parasitemia (Yamamoto et al., 2019). Additionally, we demonstrated almost a linear correlation between our malaria diagnostic system and microscopic examination for the detection of percentage parasitemia (R^2^=0.99993) in the range of 0.0001–1.0% (Yamamoto et al., 2019). The limit of detection (LOD) was 10 parasites/μL (Yamamoto et al., 2019), which was much lower than that (100–200 parasites/μL) achievable by microscopy and other malaria RDTs (Bell et al., 2006; Wongsrichanalai et al., 2007). However, these results were obtained under controlled laboratory conditions using a laboratory-cultured *P. falciparum* clone that was highly distinct from the strains found in patient samples obtained from malaria-endemic regions. In this study, we first improved the diagnostic system to enable fully automated operation. Further, we evaluated whether the automated system accurately diagnosed *P. falciparum* in individuals living in Kenya, a malaria endemic area.

## Results

### Limit of detection of the 18S rRNA nested PCR

The 18S rRNA nested nPCR method was used to detect the malaria parasites. The LOD of the parasite density was determined by nPCR using a laboratory-adapted 3D7 clone at a density range of 0.0375 to 4 parasites/μL (using 2-fold dilutions in 12 different rows) (Figure 1). A probit analysis was performed to determine the density at which the parasite could be detected with 95% confidence. The analysis revealed that LOD of the nPCR was 2.56 parasites/μL.

**Figure 1.**
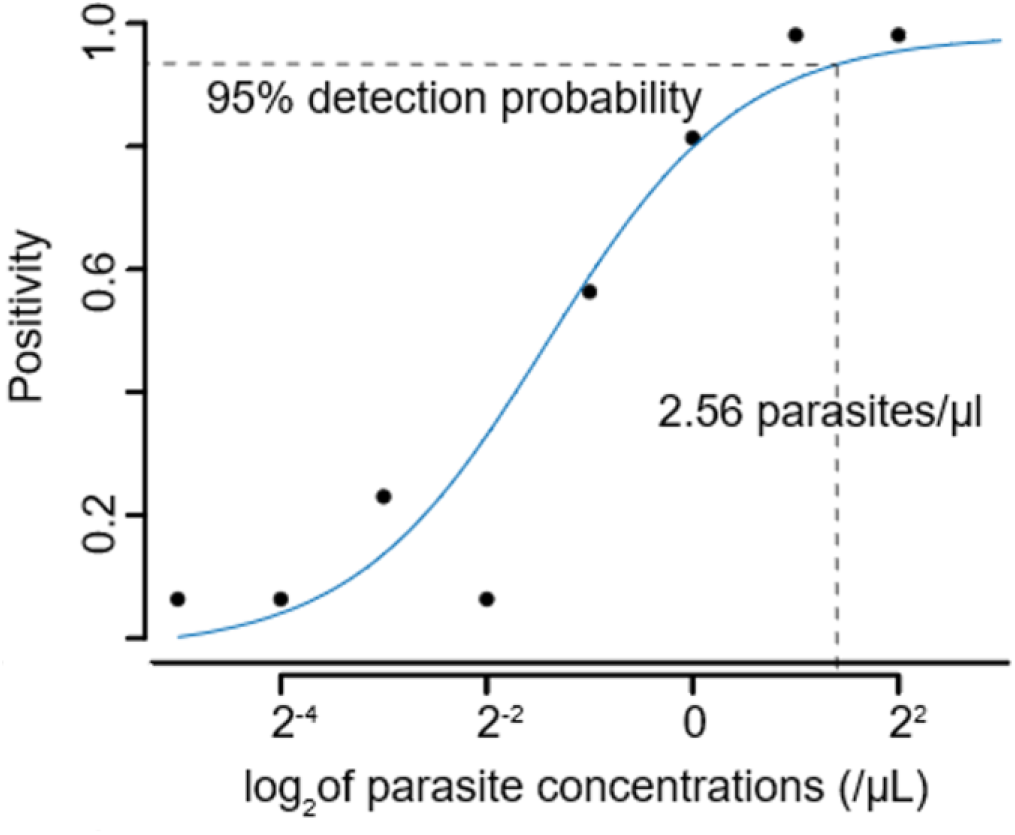
Limit of detection of 18S rRNA PCRs. 95% probability of limit of detection of 18S rRNA PCRs. Dashed lines is 95% detection probability of parasite. Continuous lines is based on the probit analysis using a serial dilution of 3D7 in vitro culture.

### Study subjects and malaria positive rates

Among the 288 school children enrolled in this study, 11 children were excluded as we did not obtain adequate amount of blood samples (n=3) or because the blood coagulated (n=8) (Figure 2). We performed nPCR analysis on the blood samples collected from the remaining 277 individuals. The analysis revealed that 56 (20.2%) samples tested positive for the malaria parasite. Among the 56 parasite-positive samples, 48 samples tested positive only for *P. falciparum*, 4 samples tested positive for *P. falciparum and P. ovale*, one sample tested positive for *P. falciparum* and *P. malariae*, and three samples tested positive only for *P. ovale* (Table 1). Since this study focused on the diagnosis of *P. falciparum*, we excluded the three samples that tested positive only for *P. ovale* from further analysis. In total, 274 individuals (95.1% of the total enrolled subjects) underwent diagnosis for malaria parasites.

**Table 1.** Demographic characteristics of 274 Kenyan individuals.

| Characteristics |  |
| --- | --- |
| Age (years) |  |
| ≤2 | 29 |
| 3–5 | 60 |
| 6–10 | 93 |
| 11≤ | 92 |
| Average | 8.4 |
| Sex |  |
| Male | 119 |
| Female | 155 |
| Hemoglobin (g/dL) |  |
| <7 | 3 |
| 7–9 | 22 |
| 10–13 | 162 |
| 13< | 87 |
| Mean (95% CI) | 12.1 ± 0.2 |
| *Plasmodium* infection status evaluated by nested PCR (number of individuals) |  |
| *Pf* | 48 |
| *Pf+Po* | 4 |
| *Pf+Pm* | 1 |
| Negative | 221 |
| Parasitemia; (%) |  |
| Median (range) | 0.04%<br>(0.00043%–1.32%) |
| *Plasmodium* infection status evaluated by RDT* (number of individuals) |  |
| Positive | 152 |
| Negative | 122 |

\*Rapid diagnostic test

**Figure 2.**
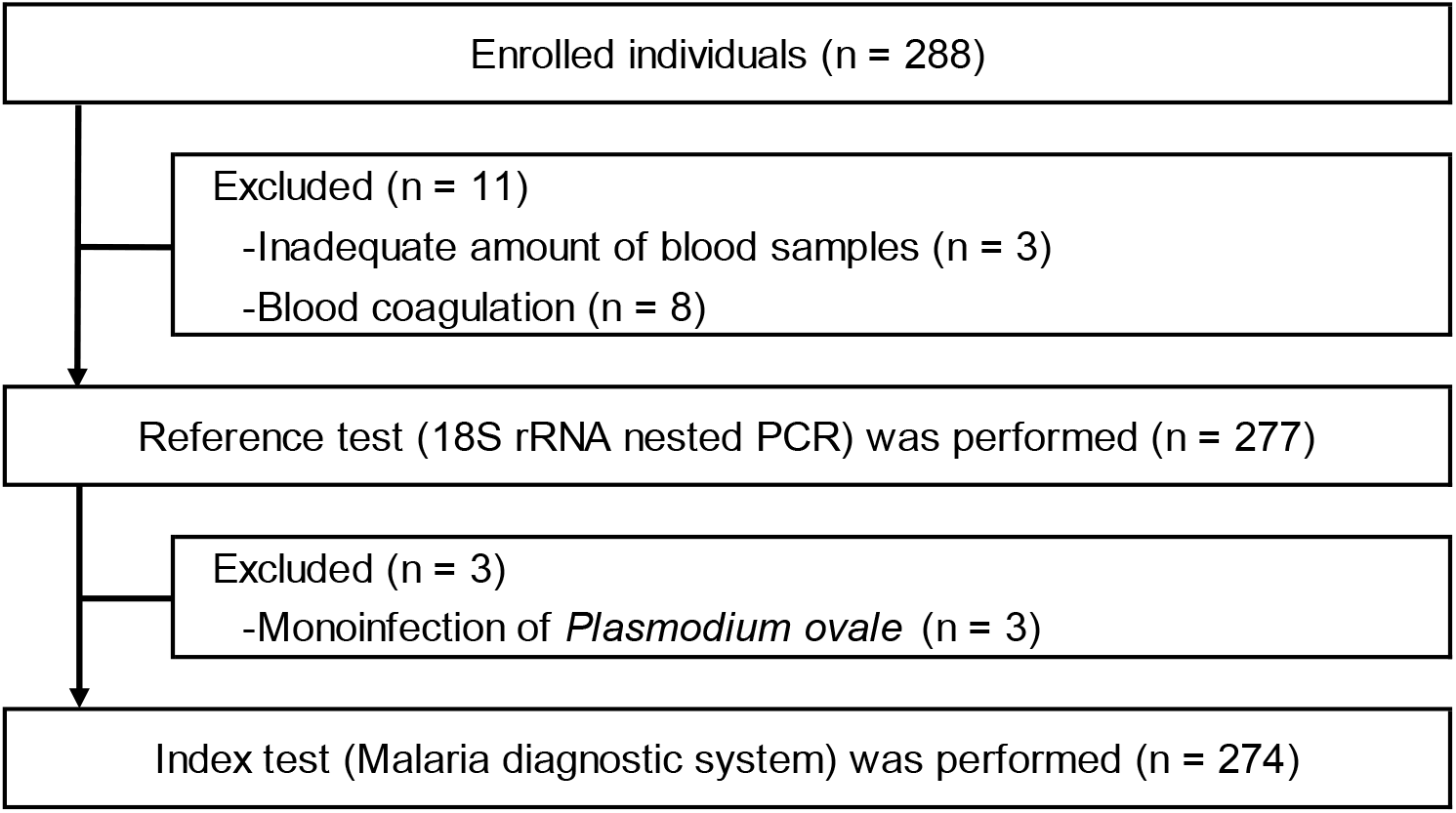
Flow diagram for the diagnosis of malaria parasites in the participants.

The mean age of the participants was 8.4 years (range: 1–16 years). Almost none of the individuals exhibited malaria symptoms (Table 1). The mean hemoglobin level among the study subjects was 12.2 g/dL. Only three individuals had a hemoglobin level of less than 7 g/dL. The median percentage parasitemia in the malaria parasite-positive samples as determined by nPCR was 0.04% (range: 0.00043–1.3%).

The sensitivity and specificity of the RDT were 98.1% and 54.8%, respectively (Table 2). The negative predictive value of the RDT was very high (99.2%), while positive predictive value was low (only 34.2%).

**Table 2.**
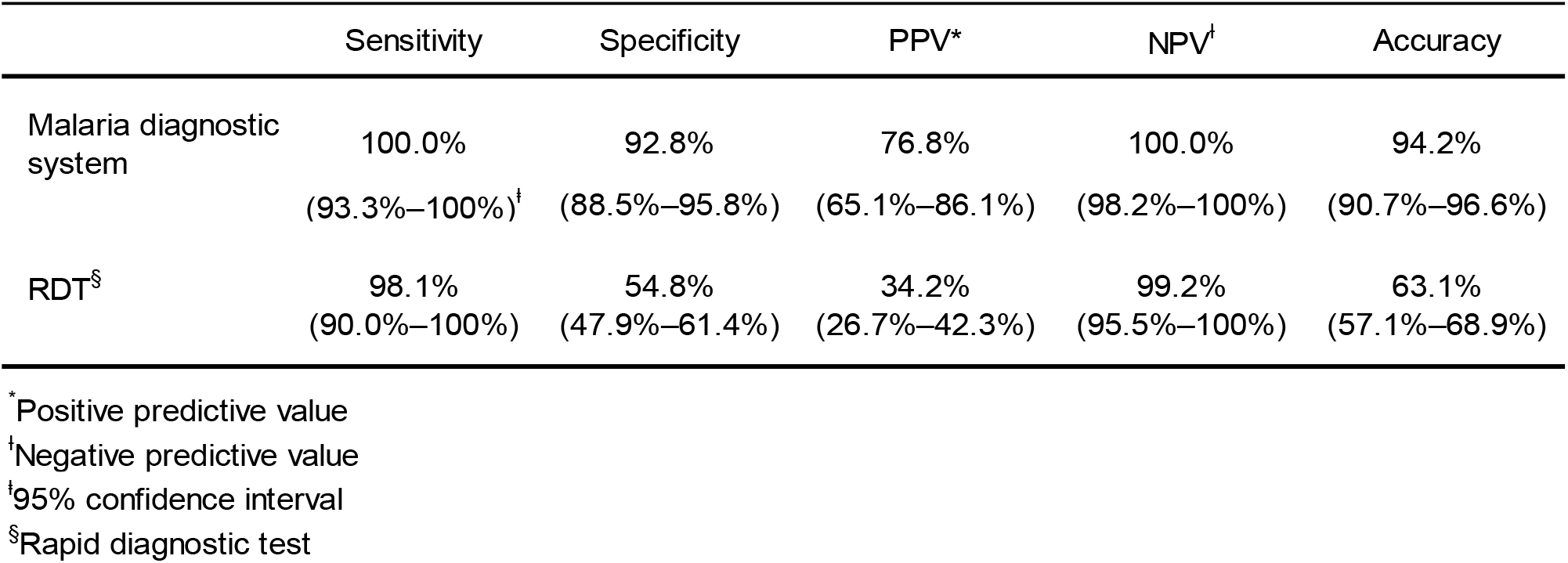
Diagnostic performance of *Plasmodium falciparum* infection in 274 Kenyan individuals.

### Development of SiO_2_ nanofiber device

Previously, we had developed a SiO_2_ nanofiber filtration system that could remove WBCs and platelets (Yatsushiro et al., 2016). By the modification of SiO_2_ nanofiber surface and strict control of pore size, only RBCs can pass through this filtration system, which can be executed without a centrifugation step. However, this SiO_2_ nanofiber filter was not incorporated within the scan disc of the diagnostic system and an additional manual step was required for the malaria diagnosis. Hence, we developed a SiO_2_ nanofiber filter that is small enough to be placed inside the scan disc. To verify the functioning of the fabricated SiO_2_, we analyzed the blood samples obtained from Kenyan individuals. We observed that the redesigned SiO_2_ nanofiber filtration system removed most of the white blood cells (WBCs). The median number of remaining WBCs on the detection area was 44 (Figure 3). This system could also efficiently remove the platelets. The mean and median percentages of the filtered platelets were 90.2% and 91.7%, respectively (Table 3).

**Table 3.** Evaluation of reformed SiO_2_ nanofiber device for the removal of platelets.

|  | Removed platelets (%) |
| --- | --- |
| Number of samples* | 11 |
| Average | 90.2 |
| Median (1st quartile, 3rd quartile) | 91.7 (83.3, 100) |

\*blood samples of healthy Japanese volunteers

**Figure 3.**
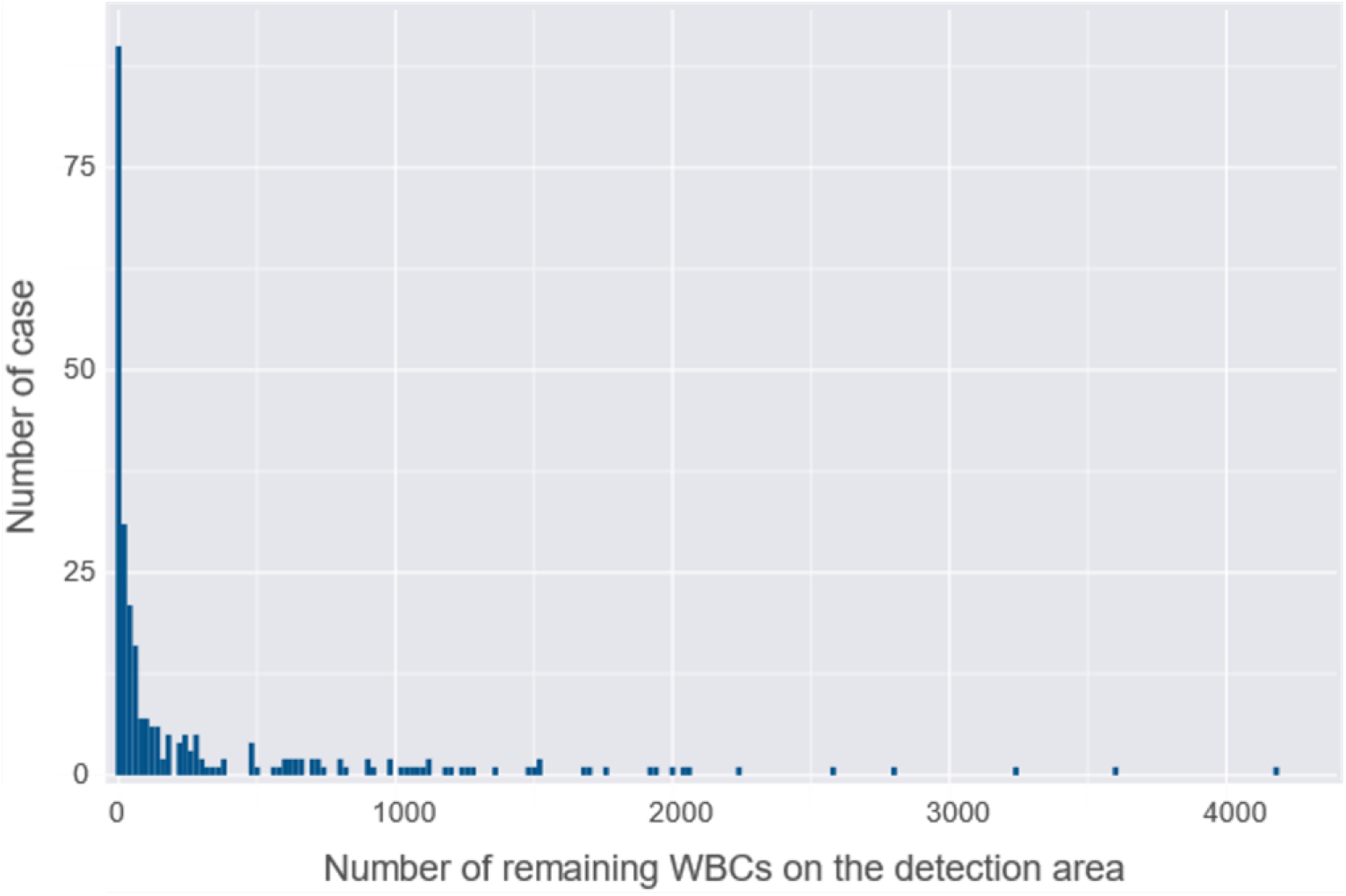
Evaluation of reformed SiO2 nanofiber device for the removal of WBCs. Remaining WBCs on the detection area. Median number of remaining WBCs was 44 (7 in 1st quartile, 280 in 3rd quartile). Blood samples from Kenya individuals were used (n = 274)

### Determination of parasitemia with the redesigned malaria diagnostic system

After the removal of WBCs and platelets by the SiO_2_ nanofiber filtration system, the RBCs were allowed to spread on the detection area, which were stained with a pre-adsorbed nuclear-specific fluorescence dye (Hoechst 34580). The staining yielded fluorescent-positive images of the malaria parasites in the infected cells (Figure 4A). Fluorescence intensity of malaria parasite was evidently lower than that of WBC (Figure 4B). Fluorescence intensities that were 1.4–2.4-times higher than those of uninfected RBCs (Figure 4B) and had a fluorescent spot with a size from 1.0 μm^2^ to 10 μm^2^ were considered to be malaria-infected RBCs {Yamamoto, 2019 #27904}. As hemoglobin has a strong absorption peak at the excitation wavelength of 400 nm (Lee et al., 2009), even RBCs can be visualized by the image reader in our diagnostic system. These images enabled us to visually count the number of RBCs and the fluorescence spots of the malaria parasites. The fluorescent image reader software identified *P. falciparum* (Figure 4B) and quantitatively measured the proportion of infected RBCs among the total counted RBCs. In general, platelets are not stained by the Hoechst 34580. However, a very small proportion of platelets were stained with this dye. In most cases, these platelets located outside RBC with distinguishable shape and fluorescent intensity from malaria parasites (Figure 4C).

**Figure 4.**
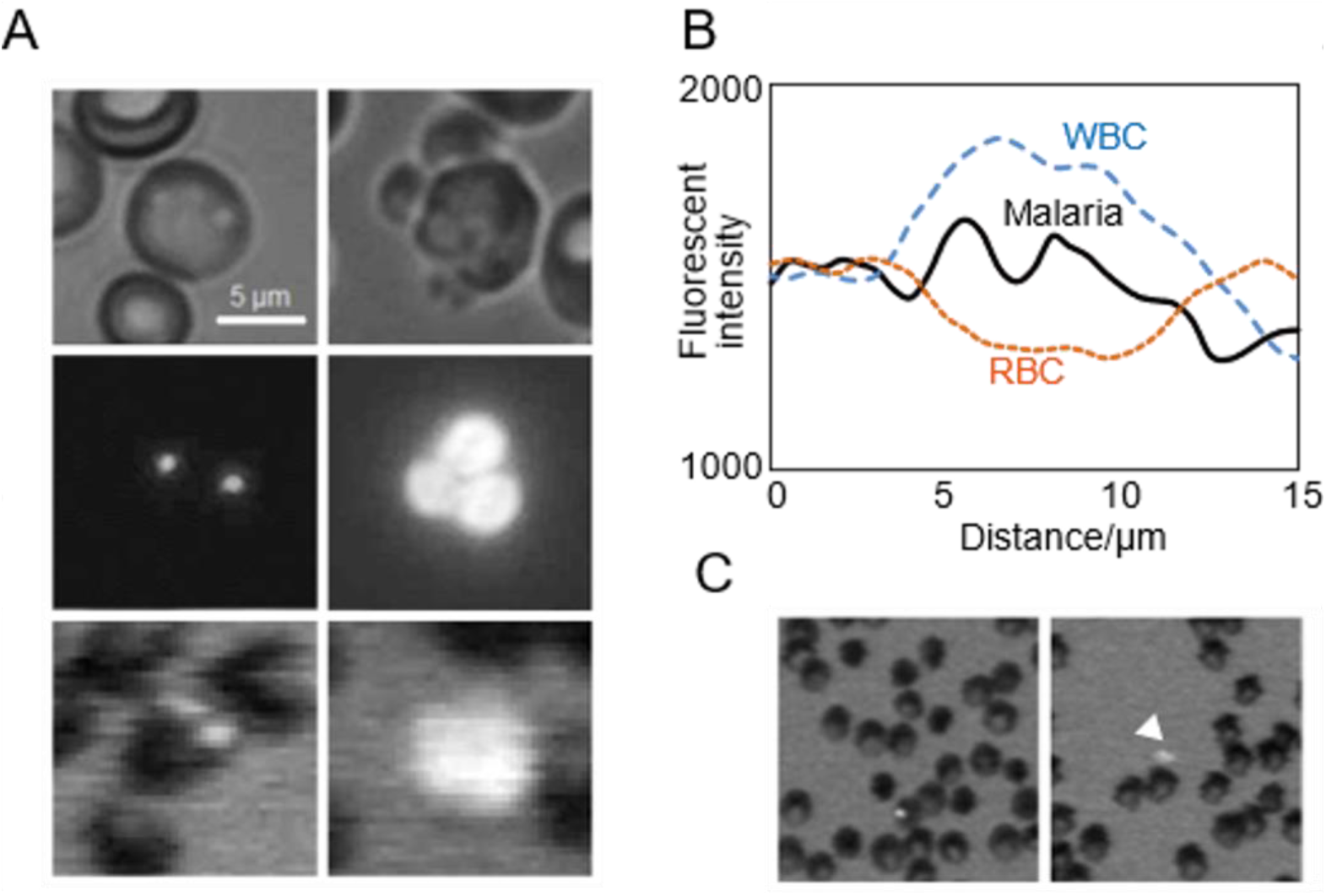
Discrimination of malaria parasites, WBCs and platelets on the fluorescent blue-ray optical system. (A) Malaria parasite (Left) and WBC dispersed (Right) on the scan disc. Differential interference-contrast microscopic images (Upper), Conventional fluorescence microscopic images (Middle) and fluorescence images by fluorescent blue-ray image reader (Lower). (B) Fluorescence-intensity profiles of RBC with malaria parasite (Black line), uninfected RBC (Orange dotted line) and WBC (Blue dotted line). The fluorescence- intensity profiles were measured along the yellow arrow in each image in Figure 4-figure supplement 1. (C) Conventional fluorescence microscopic images of malaria parasite (Left) and platelet (Right, arrowhead).

The average number of purified RBCs on the detection area was 836,863 (95% CI: 737,174–936,552) in *P. falciparum* positive samples (n = 53) and 912,556 (95% CI: 862,220–962,893) in *P. falciparum* negative samples (n = 221), respectively (Table 4). The number of fluorescent-positive spots diagnosed as malaria parasites on the detection area ranged from 23–5,130 in *P. falciparum* positive samples. However, in average 16.1 spots were also diagnosed as malaria parasites in *P. falciparum* negative samples.

**Table 4.** Number of RBCs and fluorescent-positive spots on the detection area.

| Estimates |  |
| --- | --- |
| *P. falciparum* positive samples |  |
| Number of samples | 53 |
| Mean counted RBCs (95%CI*) | 836,863 (737,174, 936,552) |
| Fluorescent-positive spots diagnosed as malaria parasites (Mean, Range) | 499 (23–5,130) |
| *P. falciparum* negative samples |  |
| Number of samples | 221 |
| Mean counted RBCs (95%CI*) | 912,556 (862,220, 962,893) |
| Fluorescent-positive spots diagnosed as malaria parasites (Mean, Range) | 16.1 (0–117) |

\*95% Confidence interval

We first determined the critical value (CV) and LOD of our diagnostic system using parasite-negative blood samples obtained from Kenyan individuals (*n*=221). The percentage parasitemia evaluated by the malaria diagnostic system for these samples ranged from 0 to 0.01% (mean: 0.0018% and standard deviation (SD): 0.0018%) (Table 5). We calculated the CV and LOD of the malaria diagnostic system from these values, which were 0.0048% and 0.0077%, respectively.

**Table 5.** Critical value (CV) and limit of detection (LOD) of parasites of the automated malaria diagnostic system.

| Estimates | % |
| --- | --- |
| Percentage of parasitemia determined by automated malaria diagnostic system in parasite negative Kenyan samples (N = 221) |  |
| Mean (%) | 0.001802 |
| SD* (%) | 0.001799 |
| Estimation with automated malaria diagnostic system |  |
| LOD (%) | 0.007722 |
| CV (%) | 0.004762 |
| Regressed to percentage parasitemia by microscopy<sup>†</sup> |  |
| LOD (%) | 0.006122 |
| CV (%) | 0.003118 |

\*Standard deviation
<sup>†</sup>Estimated LOD and CV with automated malaria diagnostic system were regressed to the microscopically determined parasitemia.

As the CV is generally used as a cutoff value for distinguishing positive results from negative results (Currie, 1995; IUPAC, 1997; Lavin et al., 2018), we adopted 0.0048% as the cutoff value for detecting parasitemia (Table 5).

The sensitivity and specificity of the malaria diagnostic system were 100% (95%CI, 93.3–100%) and 92.8% (95%CI, 88.5–95.8%), respectively (Table 2). The positive and negative predictive values were 76.8% (95%CI, 65.1–86.1%) and 100% (95%CI, 98.2–100%), respectively. The specificity obtained by our diagnostic system was significantly higher than that obtained by rapid diagnostic test (RDT) (54.8%) (P=2.2×10^−16^, McNemar’s test). We obtained false-positive results from 16 cases (Supplementary File 1), which may be due to infection from a very low number of parasites that is below the LOD of nPCR. Therefore, we verified the absence of the parasite by microscopic evaluation of 500 visual fields. Our microscopic analysis revealed that all samples that tested negative in nPCR analysis also tested negative in the microscopic analysis. There was no correlation between false-positive results and parasitemia, hemoglobin level or the number of remaining WBCs on the detection area (Table 6). However, we observed a significant correlation between age and false-positive cases (P=0.023, Welch Two Sample t-test), where false-positive cases were positively correlated with higher average age.

**Table 6.** Characteristics of false-positive cases with automated malaria diagnostic system.

|  | True positive | False positive | P |
| --- | --- | --- | --- |
| N | 53 | 16 |  |
| Age (mean, years old) | 8.4 | 11.2 | 0.02267* |
| Sex (Female%) | 43.4% | 31.3% | NS<sup>†</sup> |
| Hemoglobin (mean, g/dL) | 11.8 | 12.7 | NS* |
| Counted RBC by automated malaria diagnostic system (mean) | 836862.9 | 849898.3 | NS* |
| Remaining WBC on the detection area (mean) | 352.6981 | 253.25 | NS* |

\*Welch Two Sample t-test
<sup>†</sup>Pearson's Chi-squared test with Yates' continuity correction

To assess the variability in diagnostic accuracy across different groups of participants, we analyzed the blood samples obtained from Japanese healthy volunteers (n=40) (Table 7). The malaria parasite-positive samples were prepared by adding the *P. falciparum* laboratory clone (3D7) to the blood samples and the samples were analyzed using our diagnostic system. The analysis revealed that the diagnostic values obtained in the blood samples of Japanese healthy volunteers were similar to those obtained in the blood samples of Kenyan individuals. The sensitivity and specificity of the malaria diagnostic system were 93.3% (95%CI, 68.1 to 99.8) and 92.0% (95%CI, 74.0 to 99.0), respectively.

**Table 7.** Diagnostic accuracy of automated malaria diagnostic system for 40 Japanese volunteers

|  |  |  |
| --- | --- | --- |
| Samples |  |  |
| *P. falciparum* Positive (parasitemia)* | 15 | (0.00064-0.073%) |
| Negative | 25 |  |
| Parasitemia determined by automated malaria diagnostic system in 25 *P. falciparum* negative samples |  |  |
| Mean (%) | 0.00046 |  |
| SD (%) | 0.00024 |  |
| Estimation with automated malaria diagnostic system |  |  |
| LOD (%) | 0.00125 |  |
| CV (%) | 0.00086 |  |
| Diagnostic performance of malaria diagnostic system |  |  |
| True positive | 14 |  |
| False positive | 1 |  |
| True negative | 23 |  |
| False negative | 2 |  |
| Sensitivity (%) | 93.3 | (68.1, 99.8)<sup>§</sup> |
| Specificity (%) | 92 | (74.0, 99.0) |
| PPV<sup>†</sup> (%) | 87.5 | (61.7, 98.4) |
| NPV<sup>‡</sup> (%) | 95.8 | (78.9, 99.9) |
| Accuracy (%) | 92.5 | (79.6, 98.4) |

\*Positive samples were created by the artificial addition of *P. falciparum* laboratory clone (3D7).
<sup>†</sup>Positive predictive value
<sup>‡</sup>Negative predictive value
<sup>§</sup>95% confidence interval

### Regression analysis of parasitemia determined by revamped malaria diagnostic system and by microscopy

We evaluated the degree of correlation between the percentage parasitemia obtained by our diagnostic system and that obtained by microscopy in 53 malaria parasite-positive samples (Table 8). As the percentage parasitemia values did not exhibit a normal distribution in both methods, the data were log-transformed. Pearson’s correlation test revealed a significant correlation between the parasitemia percentage values determined by the two methods (r=0.80, P=1.2×10^−12^). A strong correlation was also obtained by Spearman’s rank-correlation test (r=0.83, P=2.6×10^−29^). These results indicated a linear correlation between the percentage parasitemia value obtained by the automated malaria diagnostic system and that obtained by microscopy.

**Table 8.** Degree of coincidence between parasitemias obtained by automated malaria diagnostic system and microscopy.

| Statistical methods | Coefficient | P |
| --- | --- | --- |
| Spearman's rank-correlation test | 0.83 | $2.6\times10^{-29}$ |
| Pearson's correlation test | 0.8 | $1.2\times10^{-12}$ |

Next, we performed a linear regression analysis to correlate the percentage parasitemia values obtained by the two detection methods (Figure 5). This analysis yielded the following equation: Predicted logarithmic transformed parasitemia by microscopy = 0.74 + (1.40 × parasitemia by automated malaria diagnostic system) with an adjusted R^2^ value of 0.6254 (P=1.13×10^−12^). As mentioned previously, the CV and LOD for our diagnostic system were 0.0048% and 0.0077%, respectively (Table 5). Using this regression equation, the corresponding CV and LOD for the microscopically determined parasitemia were estimated to be 0.0031% and 0.0061%, respectively (Table 5).

**Figure 5.**
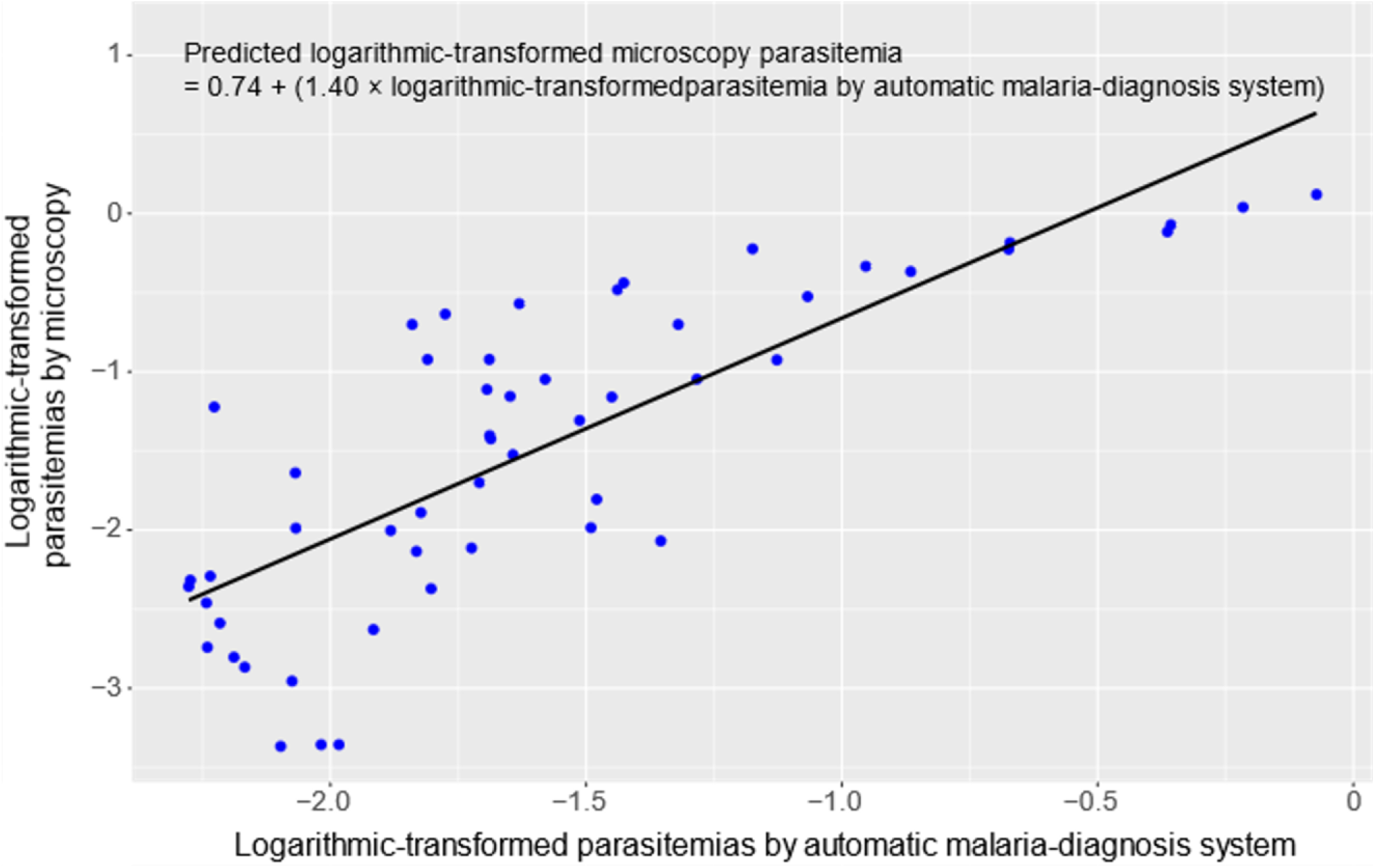
Linear regression analysis of the correlation of the percentage parasitemia obtained by the automated malaria diagnostic system with that obtained microscopically.

## Discussion

Previously, we had performed an *in vitro* evaluation of our fluorescent Blue-ray optical device-based malaria diagnostic system (Yamamoto et al., 2019). In this study, we redesigned the blood cell-filtering system and evaluated the field application of the redesigned device in a malaria-endemic region in Kenya. For this study, we enrolled Kenyan individuals who exhibited a very low parasitemia percentage (median: 0.04%) (Table 1). Our automated malaria diagnostic system could detect malaria parasites with a high sensitivity value (100%) (Table 2). Additionally, the specificity (92.8%) of our diagnostic system was much higher than that observed for other commercial RDTs (55%). The high specificity of diagnosis may prevent unnecessary administration of antimalarial drugs to non-malaria parasite-infected individuals. Consequently, this may alleviate the risks associated with the emergence and spread of drug-resistant malaria parasites (WHO, 2011).

The estimated LOD of our diagnostic system for the microscopically adjusted parasitemia in Kenyan individuals was 0.0061% (305 parasites/μL) (Table 5), which was approximately 30 times higher than that observed in our previous study (0.0002%, 10 parasites/μL) (Yamamoto et al., 2019). The LOD is mostly determined by the number of fluorescent spots that are incorrectly recognized as malaria parasites in the parasite-negative samples. One potential cause of these incorrectly detected fluorescent spots is the Howell–Jolly bodies (HJBs), which are round and small (~1 µm) nuclear remnants in the RBCs (Sears and Udden, 2012). The HJBs are morphologically similar to the malaria parasite nucleus (Lynch, 1990). Additionally, HJBs can also be stained with Hoechst 34580. These HJBs can be misdiagnosed as malaria parasites when their fluorescence intensity and size are similar to those of the parasite nucleus. In healthy individuals with a normal-functioning spleen, RBCs with HJBs are removed efficiently from the peripheral blood circulation by the spleen, and only a few of them (approximately 20/10^6^ RBCs) are present in the peripheral blood (de Porto et al., 2010). This was reported in a previous study that estimated LOD using RBCs from healthy Japanese volunteers (Yamamoto et al., 2019). However, RBCs with HJBs are not efficiently removed from the blood in patients after splenectomy or in patients with a non-functioning spleen (Davis, 1976; Pearson et al., 1969). In sickle cell disease, where splenic dysfunction begins early in life, the number of HJBs is approximately 100 times higher than that in healthy individuals (Harrod et al., 2007). Recent or acute *P. falciparum* infection was reported to induce an alteration in the splenic architecture in young children, potentially resulting in hyposplenic function (Gomez-Perez et al., 2014). Therefore, it is likely that the samples obtained in Kenya had a greater number of HJBs than the samples obtained from the healthy Japanese volunteers, which would explain the higher LOD observed in this study.

The other cause for the high LOD may be the platelets. Generally, platelets are not stained by the DNA-specific dyes. However, it is possible that a very small proportion of platelets were stained with this dye (Figure 4C). If the stained platelets attach to the surface of RBCs they could potentially be misrecognized as fluorescent spots derived from the parasitic nucleus. The blood samples used in our earlier study were obtained from the Japanese Red Cross Society and had been pretreated for RBC transfusion. Therefore, approximately 99% of the platelets had been removed before the analysis. In this study, we used fresh blood samples obtained from Kenyan individuals. To remove the platelets, we had developed SiO_2_ nanofiber filters (Yatsushiro et al., 2016), which were redesigned in this study. The revamped SiO_2_ nanofiber filters trapped approximately 90% of platelets (Table 3), which was relatively lower than the method used for the RBC purification for blood transfusions (99%). This indicated that more platelets were likely to have remained on the detection area. Further improvement of the filtration performance of the SiO_2_ nanofiber filter to decrease the number of platelets may lower the LOD. Moreover, the ability to distinguish the malaria parasites from the platelets or HJBs could be improved by enhancing the resolution of the scanned detection area. A resolution of 0.5 μm was already achieved in the current system (Yamamoto et al., 2019). However, super-resolution techniques, such as the use of a smaller spot size and an objective lens with a higher numerical aperture and/or image processing by image convolution, can potentially enhance the resolution of the detection area (Bouwhuis, 1985; J., 1988).

To reduce the number of manual steps required after blood sampling, we downsized the cell separation device and mounted it on the scan disc (Figure 6B and Figure 6-figure supplement 1). This refinement enabled the automation of the diagnostic system and reduced the amount of blood needed for diagnosis. Since the revamped separation device is made of resin by conventional injection molding, the required number of parts, manufacturing steps, and costs can be reduced.

**Figure 6.**
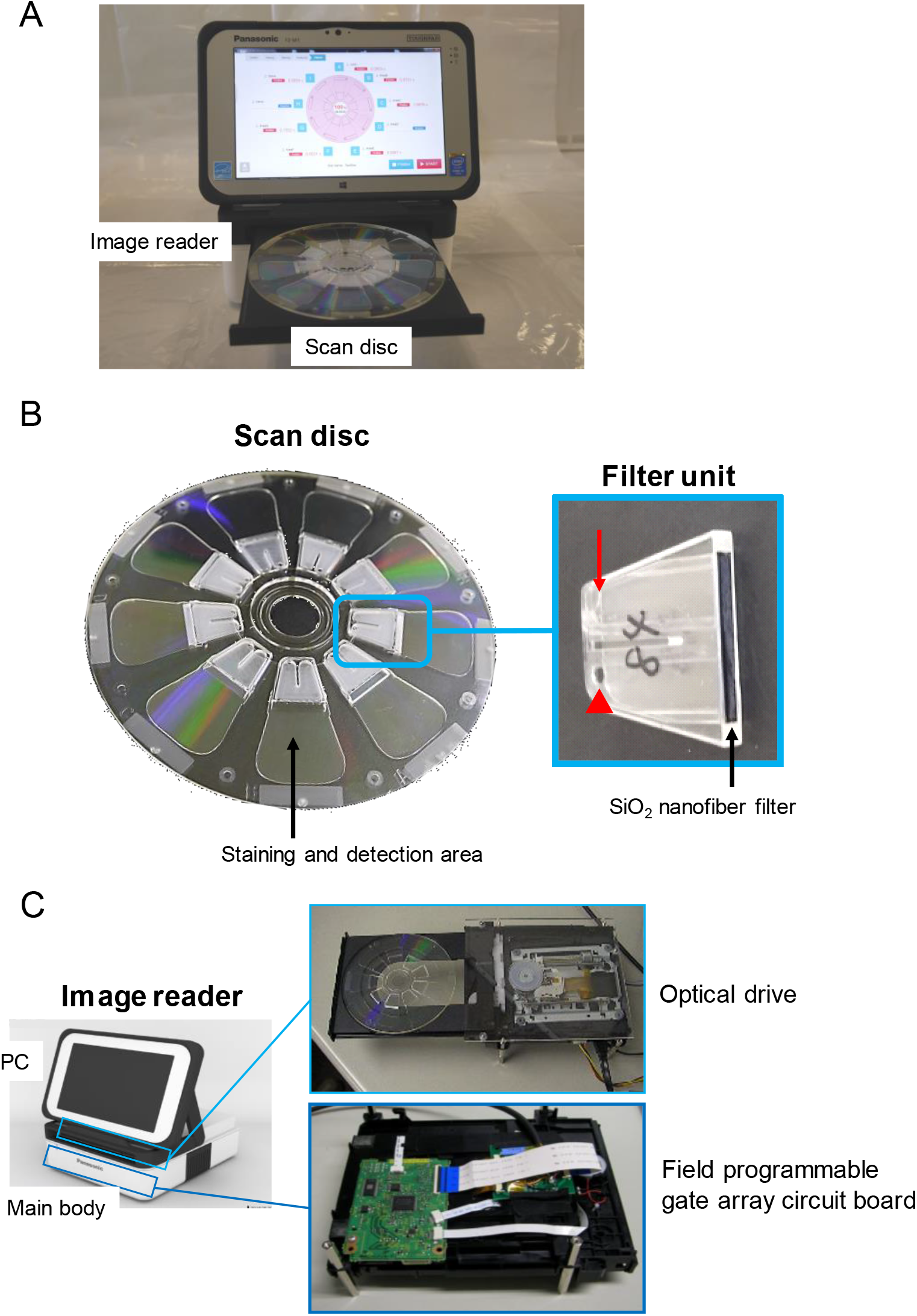
Design of the fully automated, quantitative malaria diagnostic system. (A) The fully automated, quantitative malaria diagnostic system consists of an image reader (upper) and a scan disc (lower). (B) Scan disc (left) and detailed schema of the filter unit (right). The filter unit has an air vent (arrow), and a sample injection port (arrow-head). Schematic diagram of a cross section of the scan disc is shown in Figure 6-figure supplement 1. (C) The image reader consists of a tablet PC (upper) and a main body (lower) equipped with a Blue-ray optical component.

The determination of parasitemia as a point-of-care (POC) test is extremely important as it can reduce the risk of overlooking patients with severe malaria, because high parasitemia is one of the important findings for suspected severe malaria. Furthermore, regular monitoring of the patient’s parasitemia after treatment is useful for the evaluation of therapeutic efficacy and for detecting resistance to antimalarial agents at an early stage. Various technologies have been used for the development of quantitative malaria diagnostic system, including magnetic resonance relaxation (Peng et al., 2014), flow cytometry (Tougan et al., 2018), automatic counting from digitally captured images of Giemsa-stained blood smears (Racsa et al., 2015; Rosado et al., 2017), and the evaluation of acoustic signals of vapor-generated nanobubbles from hemozoin (Lukianova-Hleb et al., 2014). However, these new devices may not be suitable for POC as they are not portable and require huge power supply. However, our malaria diagnostic system is portable, battery driven, and robust. Robustness is particularly important as malaria diagnosis is commonly performed in tropical regions with severe conditions, such as high temperature, humidity, and dusty environment. We previously reported the stability of our scan disc for several months at room temperature for the detection of *P. falciparum* (Yamamoto et al., 2019). Furthermore, our malaria diagnostic system is easy to operate and can accurately measure parasitic density independent of technical expertise. Hence, these features of our system can be advantageous for POC field use.

The effective use of POC test results is particularly important for evaluating a region’s malaria endemic status. However, the manual management of huge analog data from RDTs and microscopy is labor intensive. In our system, diagnostic data such as parasitemia and negative/positive results are digitally stored, which enables reporting of a large amount of data to the central systems such as the Ministry of Health. The effective analysis of such “big data” by sophisticated statistical methods will provide important insights into designing efficient strategies to control and/or eliminate malaria in the future.

In conclusion, we have developed an automated, quantitative malaria diagnostic system using a fluorescent Blue-ray optical device. The only manual steps involved in the use of this system are the dilution of the blood sample and its injection into a scan disc (Video 1), which would enable local health authorities to achieve the stable detection and quantification of parasitemia. Field testing of the system in Kenya revealed that the diagnostic system has a high diagnostic accuracy. These promising results indicate the potential of this diagnostic system as a valid alternative to conventional methods used at local health facilities, which lack basic infrastructure.

## Materials and Methods

### Automated malaria diagnostic system design

The automated malaria diagnostic system is primarily composed of two devices (Figure 6A): the scan disc (EZBNPC01AT, Panasonic Corp., Osaka, Japan) (Figure 6B) and the fluorescence image reader (EZBLMOH01T, Panasonic Corp.) (Figure 6C). The scan disc has a flow-path disc component and an optical disc component with a *Plasmodium* staining unit. The function of the scan disc is to isolate the RBCs and deploy them in a monolayer formation onto the staining unit. In the staining unit, malaria parasites are fluorescently stained with a nuclear-specific fluorescence stain, Hoechst 34580 (Molecular Probes Inc., Eugene, OR, USA). The fluorescence image reader detects the fluorescently stained nuclei of the malaria parasites. The identification of *P. falciparum* and the quantitative measurement of the proportion of infected RBCs among all counted RBCs are performed using a custom-made software. The system was designed for protection against particles and water based on the criteria stipulated by the International Electrotechnical Commission IP52 (IEC 60529, “Degrees of protection provided by enclosures (IP Code),” 2013). The detailed design of the diagnostic system is described in our previous study (Yamamoto et al., 2019).

### Development of SiO_2_ nanofiber device

We developed a SiO_2_ nanofiber filter that was small enough (15 mm × 2 mm, 250 µm thickness) to be placed inside the scan disc. The redesigned SiO_2_ nanofiber filter was positioned next to the blood sample injection site (Figure 6B). When the blood sample is applied to the scan disc, it first passes through the SiO_2_ nanofiber filter, where the WBCs and platelets are trapped resulting in effective RBC isolation. To develop the SiO_2_ nanofiber filter-containing scan disc, an air vent was needed to inject the 200-µL volume of sample using a pipette. However, upon injection of sample only the area between the air vent and sample injection port was filled. Therefore, to fill the entire filter unit with the sample, it was necessary to place two holes at the most distant positions apart, such that the sample injection port would be on the inner peripheral side and the air vent would be on the outer peripheral side. However, when centrifugal force was applied, the sample leaked from the hole located on the outer periphery. Hence, we created a partition in the middle and the inner structure (Figure. 6B). Through this technical improvement, we were able to establish both the sample injection port and the air hole on the inner peripheral side. Blood samples obtained from individuals in Kenya were used for evaluating the removal of WBC by the redesigned SiO_2_ nanofiber filter. We also evaluated the removal of platelets from the blood samples of healthy Japanese volunteers using the redesigned SiO_2_ nanofiber filter. We used this technique to determine the platelet count as we did not have adequate laboratory facilities to accurately determine the number of platelets.

### Study site for the evaluation of the system

A field test was conducted in February 14–23 2018 to evaluate the performance and field application of our diagnosis system. The area of the study region, which was in the Gembe East Sub-location in Homa Bay County (Mbita District, Nyanza Province, western Kenya, 0°28′24.06′′S, 34°19′16.82′′E), was approximately 12 km^2^ and included 14 villages (Minakawa et al., 2015). In this site, the annual rainfall ranges from 700 to 1200 mm, with two rainy seasons (from March to June and from November to December). All *Plasmodium* species that cause malaria in humans, except *Plasmodium vivax*, was reported in this region. Additionally, *P. falciparum* (>90%) was reported to be the most prevalent species in this region (Idris et al., 2016). The prevalence rate of malarial parasites in the study region evaluated by microscopic diagnosis and PCR was approximately 15–24% and 30–44%, respectively (Idris et al., 2016). *Anopheles gambiae sensu stricto*, *Anopheles arabiensis*, and *Anopheles funestus* are the main malaria vectors in the study region (Minakawa et al., 2002; Zhou et al., 2004).

We obtained ethical approval to conduct the study from the Kenya Medical Research Institute Ethical Review Committee (KEMRI/RES/7/3/1, SSC No. 3168), and the National Institute of Advanced Industrial Science and Technology (AIST) ethics committee (No. 2017-156).

### Blood collection

The minimum number of subjects to be enrolled for the study was determined based on the table of power estimates reported by Flahault et al (Flahault et al., 2005). According to their study, 50 infected individuals are sufficient to detect *P. falciparum* with an expected sensitivity of 95% and a lower bound 95% confidence interval value of 80%. We assumed that the prevalence of *P. falciparum* infection in the study area was 20%, which estimated that 250 individuals must be recruited for the study. This estimate is well matched to the sample size used in our study.

The study was conducted using a community-based cross-sectional survey in all villages and 14 primary/secondary schools. We collected the blood samples from school children aged 1-16 years using finger-prick technique. The consent to participate in this study was obtained directly from the children in the presence of legal guardians or through their parents. Blood collection was performed from 8 am to 1 pm. The blood samples were stored in BD Microtainer Tubes containing K_2_EDTA (Becton, Dickinson and Company, Franklin Lakes, NJ, USA) and immediately transferred to the central laboratories at the International Centre of Insect Physiology and Ecology (Nairobi, Kenya). We analyzed a maximum of 40 samples in one day. The hemoglobin level was measured with a HemoCue Hb201+ system (HemoCue AB, Ängelholm, Sweden). *P. falciparum* infection was screened using a commercial RDT kit (Paracheck-*Pf*® Rapid Test for *P. falciparum*, ver. 3, Orchid Biomedical Systems, Verna, Goa, India). We prepared thin and thick blood smears on site. The thin blood smears were fixed with methanol. All smears were stained with 10% Giemsa solution for 10 min and examined under oil immersion under a light microscope (Olympus, Co., Ltd., Tokyo, Japan) at 1000X magnification.

For molecular analysis, whole-blood samples (20 µL) were transferred onto Whatman FTA® microcards (GE Healthcare, Chicago, IL, USA). The samples were allowed to dry at room temperature and stored separately in plastic bags at −20°C. DNA was extracted from the 5.5-mm-diameter blood spots using the QIAamp DNA Micro Kit (QIAGEN, Venlo, Netherlands). The final elution volume was 20 µL. Individuals who tested positive for malaria were treated with Artemether–lumefantrine.

The purpose and procedure of the study were informed to the participants through local interpreters. Written informed consent was obtained from their parents or legal guardians.

### Diagnosis of malaria by 18S rRNA nested PCR and microscopic examination

Species-specific nPCR was used to detect the *P. falciparum* infection, as described previously (Johnston et al., 2006; Snounou et al., 1993). This method targets the 18S rRNA gene of *P. falciparum* (Johnston et al., 2006; Snounou et al., 1993). The genome of *P. falciparum* has 5–8 copies of 18S rRNA. The 18S rRNA is commonly used for DNA-based malaria detection methods (Mercereau-Puijalon et al., 2002). The reported LOD of this method varies from 1 to 50 parasites/μL (Li et al., 2014; Snounou et al., 1993; Wang et al., 2014; WHO, 2014). In this study, we determined the LOD of parasite density using a laboratory-adapted 3D7 clone at a density range of 0.0375–4 parasites/μL (using 2-fold dilutions in 12 different rows). Probit analysis was performed to determine the minimal density at which the parasite would be detected with 95% confidence. Parasitemia in *P. falciparum*-positive cases was determined by counting 10,000 RBC using thin blood smears or counting 500 WBCs using thick blood smears, while that in *P. falciparum*-negative cases was determined by counting 500 WBCs in thick blood smears through microscopic examination.

### Malaria diagnosis by the automated malaria diagnostic system

The measurement of parasitemia (percentage of parasite-infected RBCs) in the automated malaria diagnostic system comprises six steps (Figure 7A and Video 1). The first three steps are manually performed by the technician and the last three steps are automated in the device. The steps involved in the diagnosis are as follows: Step 1, the finger-prick blood sample is collected into a capillary tube. The blood (2 µL) is then manually diluted (1:100) using a buffer solution (198 µL); Step 2, the diluted blood sample is injected into the scan disc; Step 3, the scan disc is set up in the fluorescence image reader; Step 4, the diluted sample is automatically passed through the SiO_2_ nanofiber filter by centrifugal force. The purified RBCs are deployed onto the detection area in a monolayer formation (Figure 7B); Step 5, malaria parasites present in the sample are fluorescently stained (Figure 7C) and the fluorescent signals are captured by the fluorescence image reader; Step 6, the number of RBCs and malaria parasites present in the RBCs is automatically estimated from the fluorescent image by the built-in image-processing software program (EZBLMOS01T-A, Panasonic Corp.). The scan disc can analyze nine samples simultaneously in approximately 40 min. The time required for the measurement is proportional to the scanning distance of the radial direction of the scan disc. If the number of RBCs to be measured can be reduced, it is possible to shorten the measurement time.

**Figure 7.**
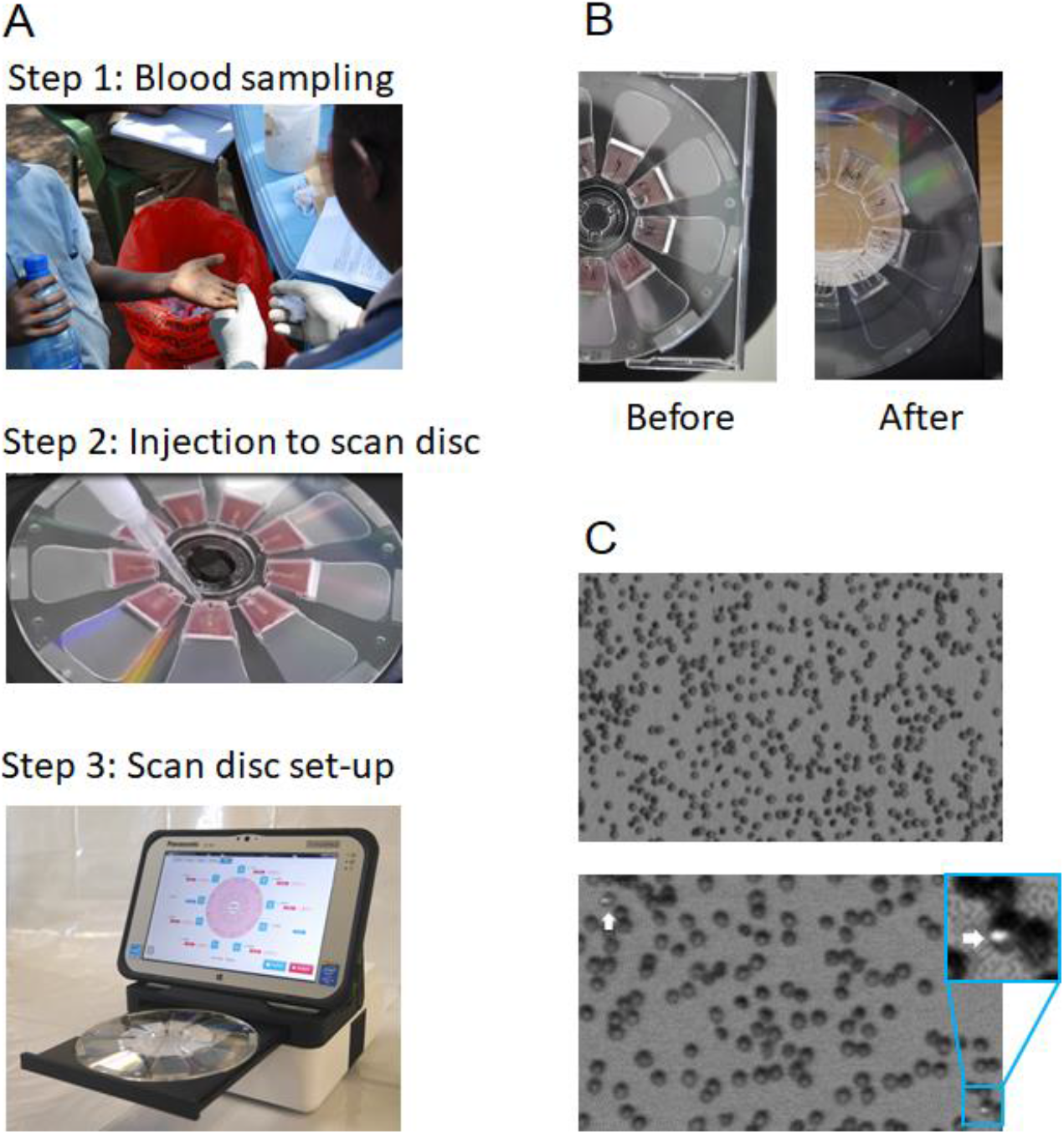
Process for malaria diagnosis using the automated malaria diagnostic system. (A) The manual steps involved in automatic malaria diagnostic system: blood sampling, injection into the scan disc, and scan disc setup. (B) Diluted samples before and after filtration by centrifugal force. (C) Fluorescent images of red blood cells (RBCs) in the detection area captured by the automated malaria diagnostic system. (Upper) RBCs are deployed in a monolayer formation. (Lower) Malaria parasites (arrows) are fluorescently stained in the detection area. High-magnification fluorescent image of *Plasmodium falciparum*-infected RBCs on the disc. The target malaria parasites were analyzed quantitatively at the single-cell level.

### Statistical analysis

The CV, which is defined as the value that produces an error probability of 0.05 when true negative samples are measured, was used as the cutoff level to distinguish the parasite-positive samples from the parasite-negative samples by this system (Currie, 1995; IUPAC, 1997). The CV was determined based on the definition provided by International Union of Pure and Applied Chemistry (IUPAC, 1997): CV = mean + 1.645 standard deviation (SD). The LOD, which is defined as the value that produces an error probability of 0.05 when samples having a LOD level are measured, was calculated using the following formula: LOD = mean + 3.29 SD (Currie, 1995; IUPAC, 1997). In this study, the samples that tested negative for the parasites in nPCR and verified by microscopic evaluation were used as true negatives for determining the CV and LOD. When the parasitemia percentage determined by our automated system was higher than the CV, the sample was considered as parasite-positive. Conversely, when the parasitemia percentage determined by the system was lower than the CV, the sample was considered as parasite negative.

The significance of discordance was measured by Welch’s t test, Chi-squared test or McNemar’s test. All statistical analyses were performed using R version 3.6.0. Exact 95% confidence intervals (CIs) were computed for sensitivity, specificity, positive and negative predictive values, and accuracy using binomial distributions with Clopper-Pearson method (Clopper and Pearson, 1934). Pearson’s correlation test and Spearman’s rank-correlation test were used to evaluate the degree of correlation between the percentage parasitemia obtained by our diagnostic system and that obtained by microscopy. A linear regression analysis was also performed to correlate the percentage parasitemia values obtained by these two detection methods. (Team, 2014)The difference was considered statistically significant when the P-value was less than 0.05.

## Acknowledgments

We thank Izumi Shibata, Satoko Fushimi and Shin-Ichiro Tachibana for technical assistance. And we thank Noriaki Terahara, Fumitomo Yamasaki, Masumi Ogawara, Takeshi Ohmori, Tsutomu Kawanishi, and Yasuhiro Mamiya for their help in developing our diagnosis system. This study was conducted at the Kenya Research Center, Institute of Tropical Medicine, Nagasaki University, Japan. This paper was published with the permission of the Director of Kenya Medical Research Institute.

---

**Figure 4-figure supplement 1.**
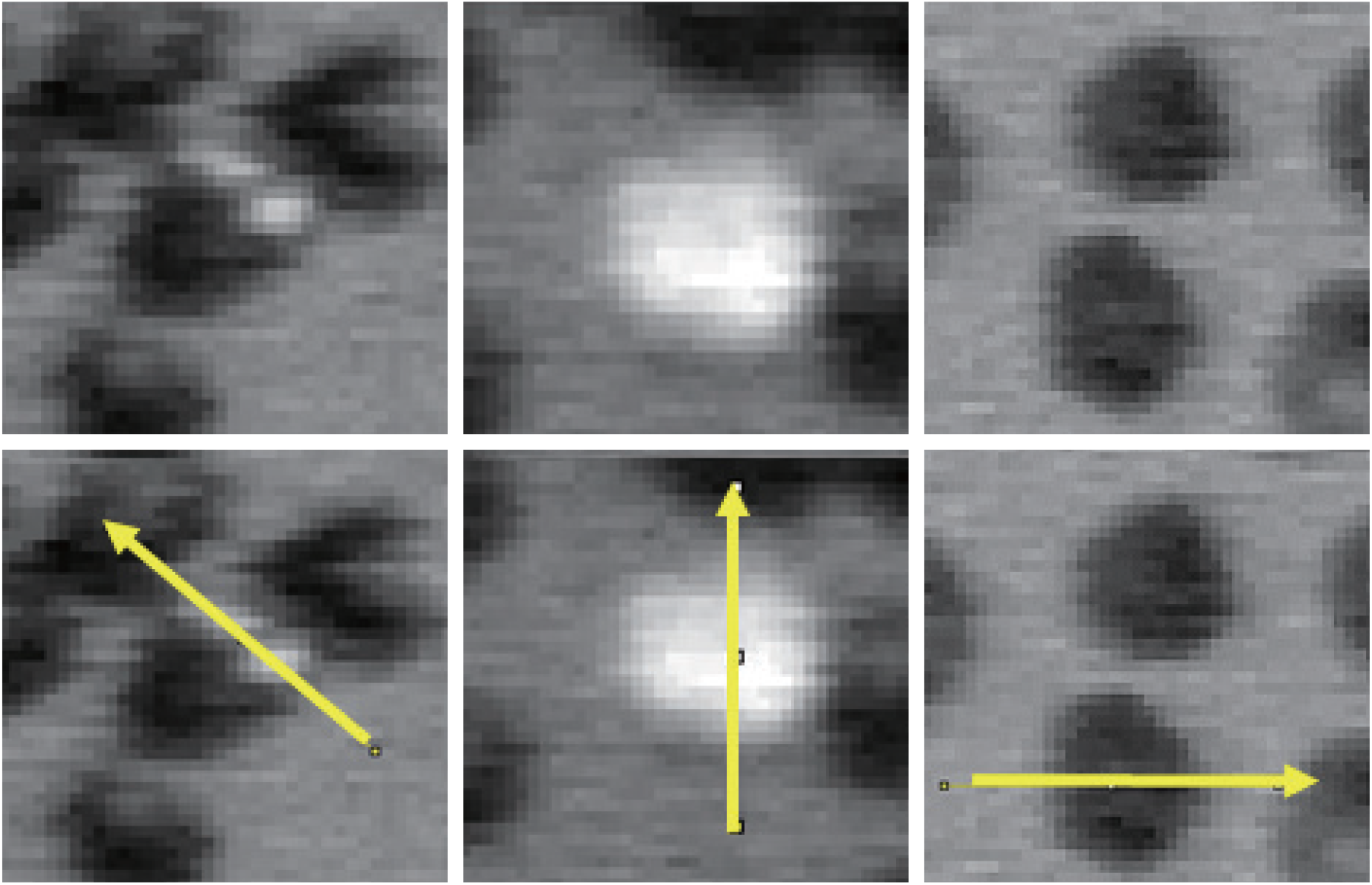
Tracing directions of fluorescence- intensity profile

**Figure 6-figure supplement 1.**
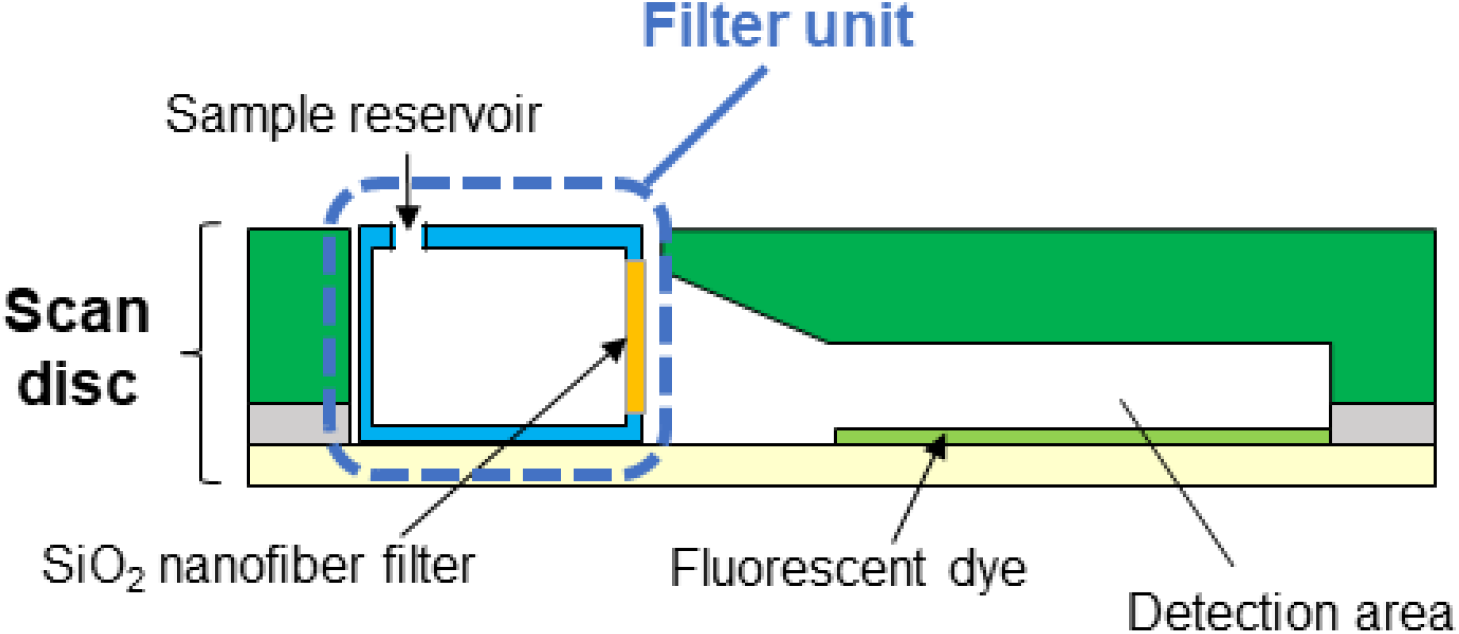
Schematic diagram of a cross section of the scan disc

Video 1. Diagnostic steps in the automated malaria diagnostic system.

Supplementary File 1. False-positive cases with automated malaria diagnostic system.

